# An epigenetic molluscicide

**DOI:** 10.1101/2023.03.21.533666

**Authors:** Nelia Luviano, Ludovic Halby, Corinne Jallet, Paola B. Arimondo, Celine Cosseau, Christoph Grunau

## Abstract

*Biomphalaria glabrata* is a fresh-water mollusk that serves as obligatory intermediate host to *Schistosoma mansoni*, agent of the neglected tropical disease schistosomiasis that affects roughly 250 Mio people. One of the ways to control the pathogenic agent is to interrupt the life cycle by eliminating the intermediate snail host though foal treatment of water bodies with molluscicides. Currently recommended molluscicides were developed in the 1950ths and lack sufficient specificity, e.g., they are toxic to fish. To provide new lead compounds for the development of a new type of molluscicides we used a rational approach based on the hypotheses that interfering with an important epigenetic mark, DNA methylation, would impede development of the snail host. We present here the compound 29, analogues-based compound that mimic substrates of DNA methyltransferases. We show that compound 29 has (i) low cytotoxicity for human cells, (ii) it inhibits DNA methylation, and (iii) it decreases fecundity in *B.glabrata*. It is therefore conceivable to produce compounds that act as specific epigenetic molluscicides.

## Introduction

Schistosomiasis is a debilitating disease of human, livestock and wild animals that affects more than 250 Mio people and an unknown, but very large number of animals. It is caused by different species of schistosome parasites that have a complex life cycle requiring asexual reproduction in different species of freshwater snails as obligatory intermediate host. Originally, schistosomes were restricted to the tropics, but since at least 2014 a hybrid of *S.bovis* and *S.haematobium* is now also endemic in Europe (Corsica, France) (Berry et al., 2014). WHO has recently redefined the strategy for schistosoma control that recommends now integrated measures including focal elimination of snail hosts (Lo et al., 2022). Molluscicides have been used successfully since the 1950^th^ to control schistosoma transmission. However, the currently available products such as niclosamide (2-amino ethanol salt of 2’, 5’-dichloro-4’-nitro salicylanilide; Bayluscide®, Bayer, Germany) are toxic for fish and not selective against vector snails arguing for the development of more selective agents (Coelho and Caldeira, 2016). It was introduced in 1956-1959 as Bayer compound 73 (GOENNERT, 1961).

One reason for the capacity of schistosoma vector snails such as *Biomphalaria glabrata* to escape different control measures is their reproduction potential: a single snail can produce 10 million descendants within 3 months (Coelho and Caldeira, 2016). We reasoned that the development of new lead compounds for molluscicides should not only kill most snails but also impede the reproduction of the survivors, in addition to be specific to mollusks, or ideally only to vector snails. A biological feature that is important for phenotypic integrity and reproduction success is epigenetic inheritance. We therefore hypothesized that impeding the epigenetic information in vector snails would decrease their viability and fecundity. Therefore, the objective of our study was to find proof of concept for a new type of epigenetic molluscicides that are selective for snails. We define “epigenetic” here as mitotically and/or meiotically heritable changes in gene function that cannot be explained by changes in DNA sequence and that impact developmental mechanisms (above the level of DNA sequence) at the origin of the phenotype and its modification across evolution (Nicoglou and Merlin, 2017). According to the WHO guidelines (https://www.who.int/publications/i/item/9789241515405), we used the South American vector snail *B.glabrata* as biological model and targeted its DNA methylation as an important bearer of epigenetic information. As in all animals, in *B.glabrata* DNA methylation is restricted to position 5 of cytosines (5-methyl-cytosine (5mC)) at the dinucleotide CpG (Luviano et al., 2021). It is catalyzed by DNA methyltransferases (DNMT).

Homology searches identified two DNMT genes in *B.glabrata: Bg*DNMT1, which is presumably the maintenance DNMT and *Bg* DNMT2, which’s function is elusive (Geyer et al., 2017). Transcription of *Bg*DNMT1 is highly abundant in the snails ovotestis (Geyer et al., 2017) enforcing the notion that DNA methylation is essential for reproduction. Accordingly, we and others had previously shown that pharmacological inhibition of DNMT activity results in phenotypic changes and reduced reproduction rate (Luviano et al., 2021). Aberrant DNA methylation is a hallmark of cancer and human DNMTs have been targets for the development epidrugs against the disease for many years.

We had previously designed and synthetize a first chemical library of bisubstrate analogues-based compounds that mimic both substrates of DNMT: S-adenosyl-L-methionine (SAM) and the deoxycytidine and block the catalytic pocket of of human DNMT3A and DNMT1 (Halby et al., 2017).

We identified among those compounds that were able to induce gene promoter demethylation associated with gene reactivation in different cancer cell lines while others were inactive. We wondered whether in this chemical library there would be some compounds active only on the *B.glabrata* DNA methylation.

After a first-pass survey of previously tested molecules compound **29** (Figure 1) emerged as a suitable candidate, as it resulted completely inactive against human DNMT3A and DNMT1 (Halby et al., 2017).

**Figure 1:**
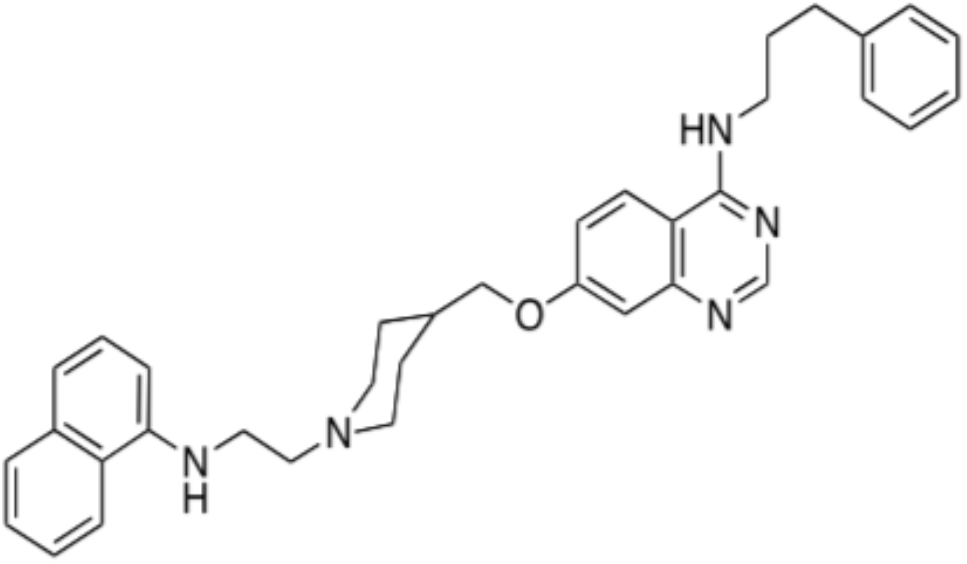
Chemical formula of compound **29**; (Halby et al., 2017)

Here we provide evidence that compound **29** is (i) little cytotoxic for human cells, (ii) it inhibits DNA methylation in *B.glabrata*, and (iii) it decreases fecundity in *B.glabrata*. It is therefore conceivable to produce compounds that act as specific epigenetic molluscicides.

## Results

### Compound 29 shows low toxicity for mammalian cells

The human liver cancer cell line HepG2 and the human leukemia MV41-1 cell lines were used for cytotoxicity tests. The potent human DNMT3A inhibitor (IC50 1μM) compound **68** of the same chemical library was used as reference. In both cell lines compound **29** was less cytotoxic than compound **68** (Figure 2): in HepG2, EC50 of 35μM for **29** against 6.9μM for **68**, and in MV4-11 4.4μM versus 1.9μM).

**Figure 2.**
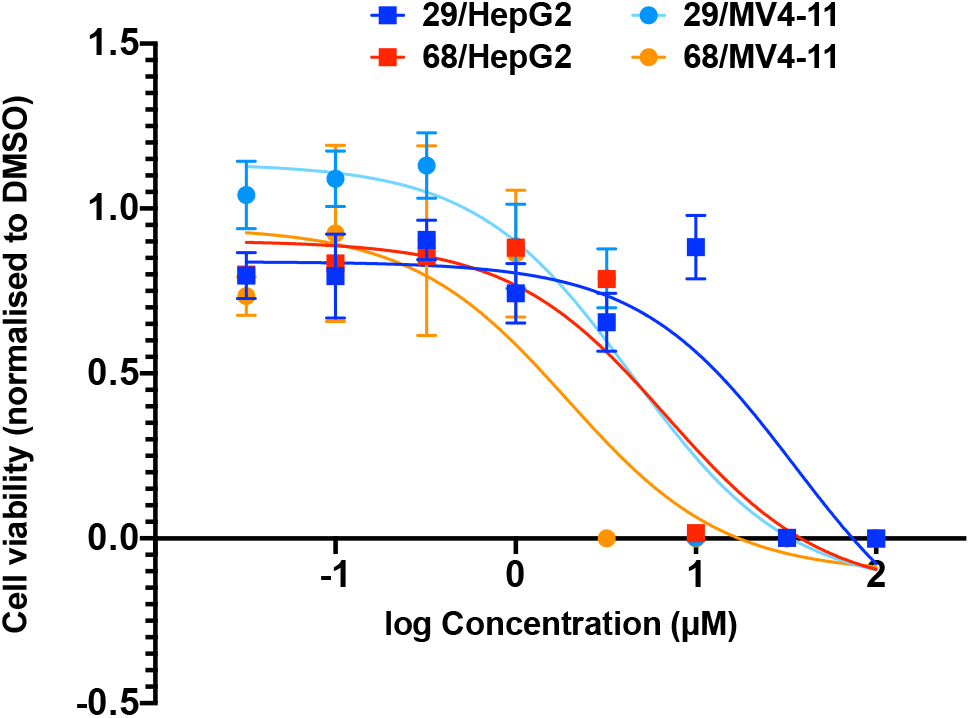
Cell viability of compounds **29** (blue) and **68** (red) compared to DMSO in HepG2 cells (squares) and MV4-11 cells (circles).

As outlined above, earlier data indicated absence of inhibitory activity against human DNMTs for compound **29** (Table 2).

**Table 1.** Summary table of inhibition of enzymatic activity and EC50 for comounds 29 and 68 from (Halby et al., 2017).

|  | % inhibition DNMT3A-c |  | % inhibition DNMT1 |  | EC<sub>50</sub> DNMT3A-c | EC<sub>50</sub> DNMT1 |
| --- | --- | --- | --- | --- | --- | --- |
|  | 32 μM | 10 μM | 100 μM | 32 μM |  |  |
| Compound **29** | 0% | 0% | 0% | 0% | >> 32 μM | >> 100 μM |
| Compound **68** | 83 % | 86 % | 69 % | 24 % | 1.1 ± 1.2 μM | 100 ± 3 μM |

### Compound 29 inhibits DNA methylation of Biomphalaria glabrata through 2 generations

Global 5mC content was measured by a quantitative dot blot technique. Results showed that the compounds **29** and **68** reduced the global 5mC% 2-fold in the F0 generation (Figure 3a, red bars). The compound **29** led to a 1.5-fold reduction of the global 5mC% in the F1 generation compared to control and compound **68** led to a 2-fold higher 5mC% (Figure 3 b, red bars). There was not significant difference between compound **29** and **68**. We also tested the DNA methylation inhibitor zebularine, as it has a different mechanism of action (Champion et al., 2010). Zebularine did not induce significant effects in 5 mC% (Figure 3 a-b, gray bars).

**Figure 3.**
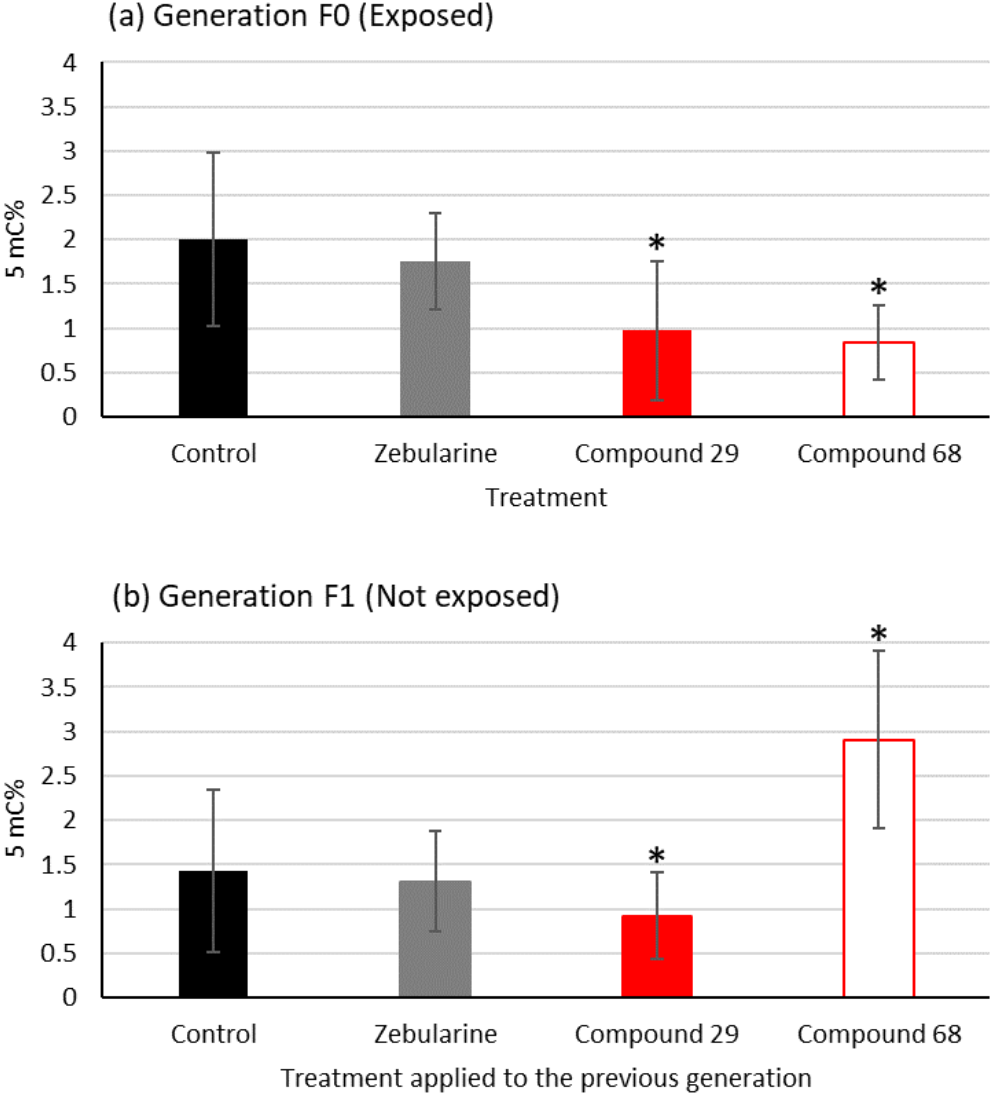
5mC % of *B. glabrata* snails upon treatments with compounds **29**, **68** and Zebularine at a concentration of 10 μM, error bars represent SD, *n* = 30 per treatment. (a) 5mC % in the F0 generation (exposed), black bar for control, gray bar for Zebularine, red bar corresponds to compound **29** and the white bar with red outline represents the compound **68.** (b) 5mC % in the non-exposed F1 generation. Compounds are the ones used in F0. Mann-Whitney Wilcoxon test was applied, if not otherwise indicated, between treatment and control significant differences are represented by *.

### Compound 29 decreases survival and fecundity in treated B. glabrata snails

Compound **29** triggered a significant mortality in snails, with a 61% survival at the end of the 70 days period (Figure 4), and significant differences were found in survival percentage between snails treated with compounds **29** and **68** compared to the control group (χ2=14.17, p=0.0002 for compound 68 and χ2=7.54, p=0.006 for compound 29).

**Figure 4.**
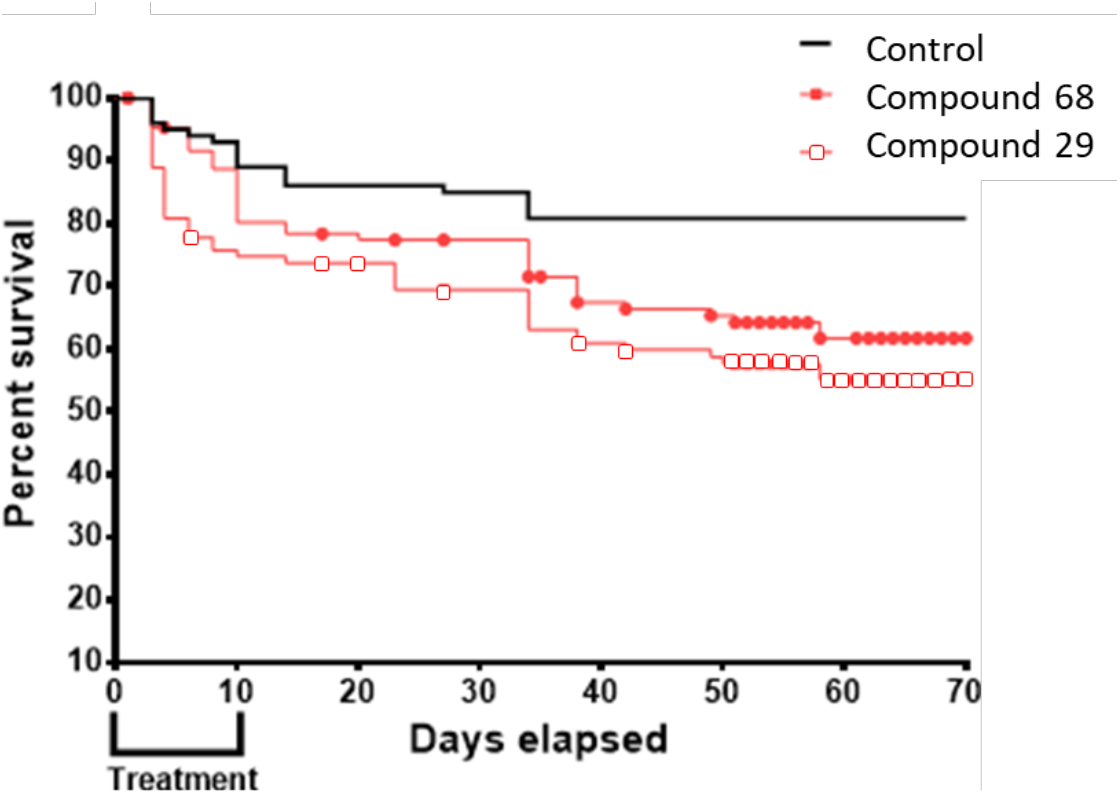
Kaplan-Meier survival curves upon treatment with compound 29 (red line with red circles) and compound **68** (red line with white squares).

Compound **29** also presented a significant difference in the number of offspring compared to control group (p=0.0002) (Table 2).

**Table 2.** Contingency table of fecundity of the snails exposed to zebularine, compounds **29** and **68**. Total number of laid eggs (first row), number of non-hatched eggs (second row) and number of offspring snails (third row). Fisher’s exact test was applied, significant differences compared to control group are marked with ** for p<0.0005.

|  | Control | zebularine | Compound 29 | Compound 68 |
| --- | --- | --- | --- | --- |
| Total number of laid eggs | 191 | 326 | 208 | 170 |
| Number of non- hatched eggs | 152 | 301 | 192 | 129 |
| Number of offspring | 39 | 25 ** | 16** | 41 |

## Discussion

Our findings indicate that the bisubstrate analogue compound **29** designed as an inhibitor of DNMTs has a significant effect on DNA methylation in the aquatic parasite vector snail *B. glabrata*. It also decreases survival and fecundity in these snails. It offers therefore the possibility to develop a new type of molluscicide. However, in order to be used in an ecological realistic context, it would be necessary to make such a compound environmentally safe i.e. other aquatic organisms, in particular vertebrates. Our cytotoxicity test indicates that this should be the case for compound **29** that is less toxic than the chemically parent compound **68**, a potent inhibitor of human DNMT3A. Indeed, compound **29** resulted inactive against human DNMT1 and DNMT3A and could thus be the starting point for a selective compound for DNA methylation in *B. glabrata*.

Interestingly, compound **29** is also inefficient against *Plasmodium falciparum* (Nardella et al., 2020) ACS Cent. Sci., 6, 16-21} lending further support to a specificity against snails.

Environmental safety and stability of compound **29** needs to be tested further and we anticipate that it needs chemical optimization before a possible development. Nevertheless, we believe that the compound has the potential as a lead product to be further improved and to serve as a base for epigenetic environmental engineering.

## Materials and Methods

### Ethics statement

*B. glabrata* albino Brazilian strain (*Bg*BRE) was used in this study. The snails were maintained at the IHPE laboratory facilities; they are kept in aquariums and fed with lettuce *ad libitum*. Housing, breeding, and animal care were done following the national ethical requirements.

### DNMTi treatment and snail phenotyping

Compound **29** and **68** were synthetized as reported in Halby *et al.* 2017 (Halby et al., 2017)and were used to treat *Biomphalaria glabrata* mature adults (8 weeks aged) with a 5-7 mm shell diameter for 10 days. Stock solutions at 10 mM were made for each compound in ultrapure Milli-Q water and added to 1L of water (final concentration 10 μM) in an aquarium with 100 snails for each compound. Another aquarium without any compound was maintained as the control group with 100 snails. The water was replaced once with fresh drug-containing water at the same concentration, the replacement was performed after 3 days and 22 h. After 10 days of exposure, the drug was removed and replaced by drug-free water. Snails were then raised in the plastic tank for 70 days, during which different life history traits were measured. Mortality was measured at days 3, 4, 6, 8 of the treatment and then each week. The egg-capsules laid by the snails of the generation F0 were separated each week to raise the F1 generation in another plastic container. The fecundity of the control and treated snail were analyzed by recording the number of eggs and the number of hatched snails, to then calculate the hatching rate (number of hatched snails/ total number of eggs ×100). At day 70, snails of the generation F0 and F1 were collected and stored wrapped in aluminum sheets individually at −20 °C.

### DNA extraction and dot-blot DNA methylation assays

The NucleoSpin® Tissue Kit (Macherey-Nagel, Düren, Germany) combined with the use of zirconia beads (BioSpec, USA, Cat. No. 11079110z) as described previously (de Lorgeril et al., 2018) was used for DNA extraction from whole body without shell of 30 *B. glabrata* snails per treatment group. The global 5mC of the 30 snails per treatment was measured by the dot blot method based on the recognition of 5 methylcytosine (5mC) by anti-5mC antibody (Abcam, Cat. No. ab73938, Lot: GR278832-3) as described previously (Luviano et al., 2018).

#### Cytotoxicity test

Human HepG2 (hepatocellular carcinoma, DSMZ) and MV4-11 (acute monocytic leukaemia, DSMZ) cells were grown as advised by the provider. Cytotoxicity was evaluated after 72 h of drug treatment in 96-well plates by ATPlite (PerkinElmer) according to the manufacturer’s protocol. Experiments were run in technical triplicates. GraphPad Prism 8 was used to interpolate IC_50_. DMSO was used as control.

## End Matter

### Author Contributions and Notes

N.L, L.H., P.B.A., C.C. and C.G. designed research, N.L. performed work on *B.glabrata*, L.H. and P.A. produced the compounds, C.J. performed cell line work, N.L and all authors wrote the paper.

The authors declare no conflict of interest.

## Acknowledgments

“This study is set within the framework of the « Laboratoire d’Excellence (LabEx) » TULIP (ANR-10-LABX-41) and CeMEB (ANR-10-LABX-04-01). N.L. was supported by a PhD grant of the Region Occitanie, France and the UPVD.

## References

Berry, A., Moné, H., Iriart, X. et al. 2014. Schistosomiasis Haematobium, Corsica, France. Emerging Infectious Diseases, 20,

Champion, C., Guianvarc’h, D., Sénamaud-Beaufort, C., Jurkowska, R.Z., Jeltsch, A., Ponger, L., Arimondo, P.B. and Guieysse-Peugeot, A.-L. 2010. Mechanistic Insights on the Inhibition of C5 DNA Methyltransferases by Zebularine. PLOS ONE, 5, e12388

Coelho, P.M.Z. and Caldeira, R.L. 2016. Critical analysis of molluscicide application in schistosomiasis control programs in Brazil. Infectious Diseases of Poverty, 5, 57

de Lorgeril, J., Lucasson, A., Petton, B. et al. 2018. Immune-suppression by OsHV-1 viral infection causes fatal bacteraemia in Pacific oysters. Nature Communications, 9, 4215

Geyer, K.K., Niazi, U.H., Duval, D. et al. 2017. The Biomphalaria glabrata DNA methylation machinery displays spatial tissue expression, is differentially active in distinct snail populations and is modulated by interactions with Schistosoma mansoni. PLOS Neglected Tropical Diseases, 11, e0005246

Goennert, R. 1961. Results of laboratory and field trials with the molluscicide Bayer 73. Bull World Health Organ, 25, 483–501

Halby, L., Menon, Y., Rilova, E. et al. 2017. Rational Design of Bisubstrate-Type Analogues as Inhibitors of DNA Methyltransferases in Cancer Cells. J Med Chem, 60, 4665–4679

Lo, N.C., Bezerra, F.S.M., Colley, D.G. et al. 2022. Review of 2022 WHO guidelines on the control and elimination of schistosomiasis. Lancet Infect Dis, S1473-3099(22)00221

Luviano, N., Diaz-Palma, S., Cosseau, C. and Grunau, C. 2018. A simple Dot Blot Assay for population scale screening of DNA methylation. bioRxiv,

Luviano, N., Lopez, M., Gawehns, F., Chaparro, C., Arimondo, P.B., Ivanovic, S., David, P., Verhoeven, K., Cosseau, C. and Grunau, C. 2021. The methylome of Biomphalaria glabrata and other mollusks: enduring modification of epigenetic landscape and phenotypic traits by a new DNA methylation inhibitor. Epigenetics & Chromatin, 14, 48

Nardella, F., Halby, L., Hammam, E. et al. 2020. DNA Methylation Bisubstrate Inhibitors Are Fast-Acting Drugs Active against Artemisinin-Resistant Plasmodium falciparum Parasites. ACS Central Science ACS Cent. Sci., 6, 16–21

Nicoglou, A. and Merlin, F. 2017. Epigenetics: A way to bridge the gap between biological fields. Studies in History and Philosophy of Science Part C: Studies in History and Philosophy of Biological and Biomedical Sciences,

